# Restoration of the Lost Human Beta Defensin (hBD-1) in Cancer as a Strategy to Improve the Efficacy of Chemotherapy

**DOI:** 10.1101/2023.04.03.535411

**Authors:** Raghu S. Pandurangi, Thillai V. Sekar, Ramasamy Paulmurugan

## Abstract

Both innate and adaptive immunity are the important components of the human defense system against various diseases including cancer. Human Beta Defensin (hBD-1) is one such immunomodulatory peptide which is lost at high frequencies in malignant cancers, while high levels of expression are maintained in benign regions making it a potential biomarker for the onset and metastasis of the disease. Loss of putative function of hBD-1 as a tumor suppressor gene combined with the defects in apoptosis pathways (CD95, ASK1) make tumor cells insensitive to chemotherapy and render it ineffective.

Triple negative breast cancer (TNBC) is an aggressive form of breast cancer for which no targeted therapy works due to lack of biomarkers (ER, PR and HER2 negative). That makes chemotherapy as a first line of treatment despite high side effects. TNBC is known for avoiding immunosurveillance and desensitizing themselves to intervention by dysregulating cell death pathways (CD95 & ASK1) and developing resistance to chemotherapy A priori Activation of Apoptosis Pathways of Tumor often referred to as AAAPT is a novel targeted tumor sensitizing technology which sensitizes low responsive and resistant tumor cells to evoke a better response from the current treatments for TNBC. Here, we show that hBD-1 is shown to target tumor specific biomarker Trx, activates dual cell death pathways CD95 and ASK1 (apoptosis stimulating kinase) to sensitize TNBC cells to chemotherapy drug Doxorubicin. As far as we know, this is the first-time injection of hBD-1 in TNBC mouse model to prove the restoration of hBD-1 back to the basal level can sensitize cancer cells which resulted in significant reduction of tumor volume in TNBC mouse model‘ in vivo. Sensitizing the low or non-responsive tumor cells by AAAPT and making chemotherapy work at lower doses may lead to the significant reduction of dose related side effects and may expand the therapeutic index of the current treatments.

## Introduction

Tumor cells have a remarkable ability to survive and proliferate by evading the cell death pathways^1^ and immune surveillance^2^. Loss of putative function of hBD-1 as a tumor suppressor gene, combined with defect in apoptosis pathways (e.g. CD95, ASK1) makes tumor cells insensitive to chemotherapy^3-5^. The ability of cancer cells to circumvent interventions by desensitizing themselves to interventions irrespective of the nature of intervention and avoiding immune surveillance is a serious issue in designing the treatment options for cancer patients. Novel methods are needed to understand the molecular biology of the desensitization and immune surveillance process to offset the tactics adapted by cancer cells which circumvent the interventions.

The reported loss of hBD-1 in malignant tumor compared to benign tumor, its correlation to malignancy and restoring the loss of hBD-1 potentially to achieve tumor regression in tumor models is significant in treating cancer^6^. The role of hBD-1 as a chemoattractant, defending agent for microbial infection, its inverse correlation between hBD-1 expression and invasive potential in oral squamous cell carcinoma (OSCC), prostate, renal and kidney cancer is well known^7^. Expression of hBD-1 is shown to be an independent predictor for survival of OSCC patients. Considering the existing literature^1-7^, it may be safe to claim that hBD-1 may contribute to the host antitumor immunity and if hBD-1 is lost at some point during the transition from benign to malignant tumor, the host may be less likely to recognize tumor as foreign, and may not be able to recruit the dendritic cells and memory T cells which means tumor survives. The loss of expression of this protein by transformed epithelium may therefore inhibit host immune recognition or defensin-mediated tumor cell cytotoxicity.

hBD-1 is an innate and adaptive immunity protein which is shown to a) induce the migration of immature dendritic cells and memory T cells^8^, b) trigger a type 1 immune response *in vivo* against the tumor antigen^9^ and more importantly, c) is a candidate tumor suppressor gene located on chromosome 8p23^10^. Restoration of the loss of hBD-1 back to the base level *in vitro* and *in vivo* (TNBC model)^11^ is established which led to the significant cell death and tumor regression respectively. This indicates that hBD-1 loss could be a potential biomarker for the onset of cancer which may lead to the use of hBD-1 expression as a valuable diagnostic tool to prevent the onset or monitor cancer progression.

Added to the immunity compromise through the loss of hBD-1, we have also shown that how hBD-1 targets tumor specific biomarker Trx and oxidizes Trx to activate Apoptosis Stimulating Kinase (ASK1) cell death pathway in plant defensin Def1 which is very similar to hBD-1 and resulted in sensitizing low-responsive tumor cells to chemotherapy (e.g. Doxorubicin)^12^. Consequently, IC-50 of Doxorubicin and several other front-line treatments for TNBC patients (paclitaxel, gemcitabine) is reduced by 8-10 times *in vitro* leading to the reduction of dose related toxicity in normal epithelial breast cells (MCF-10A) and particularly in cardiomyocytes. Hence, simultaneous restoration of the lost hBD-1 combined with sensitizing low-responsive tumor cells may improve the efficacy of the existing cancer treatments including chemotherapy and potentially, radiation and immunotherapy.

Here we report the synthesis of hBD-1 in large quantities using indegenously developed controlled oxidation protocols, antitumorigenic potential and synergy with first line treatment Doxorubicin for many cancers including triple negative breast cancer (TNBC). Restoration of hBD-1 with high synergic potential with chemotherapy may well be catered to personalized therapy for TNBC patients since they do not express biomarkers (e.g. ER, PR and HER2 negative) for which current receptor based targeted drugs are designed^13^. However, the loss of hBD-1 in TNBC makes it a good candidate for personalized treatment. Unfortunately, chemotherapy or cocktail of chemotherapy for TNBC patients as a stand-alone treatment may not be desirable but currently is the only option despite having high off-target toxicity^14^.

## Results

### A: Synthesis of hBD-1

hBD-1 is 36 amino acids cationic peptide with conserved folding through six cysteines forming three S-S folding. In general, hBD-1 is synthesized either by chemical method with protecting cysteine thiols and forming a folding or through a recombinant methods. Both are difficult processes yielding mostly partially folded or unfolded peptide leading the loss of antimicrobial activity. Here we, report a simple controlled oxidation method protocol which leads to > 96 % purity assessed through LC-MS and also by MS-MS. Figure 1 shows the formation of hBD-1 from linear peptide to cyclized peptide when the oxygen was slowly bubbled through solution of hBD-1 in distilled water. The all-reduced, hBD-1 was synthesized using peptide synthesizer (PSP510, ArrayIt, Sunnyvale, CA). The linear peptide was slowly oxidized by bubbling oxygen in a sealed tune for 45 hrs to form the three intramolecular S-S folding. The final peptide was purified to > 97 % purity by HPLC and characterized by MS before using it for *in vitro/in vivo* analysis which showed the biological activity.

**Fig 1:**
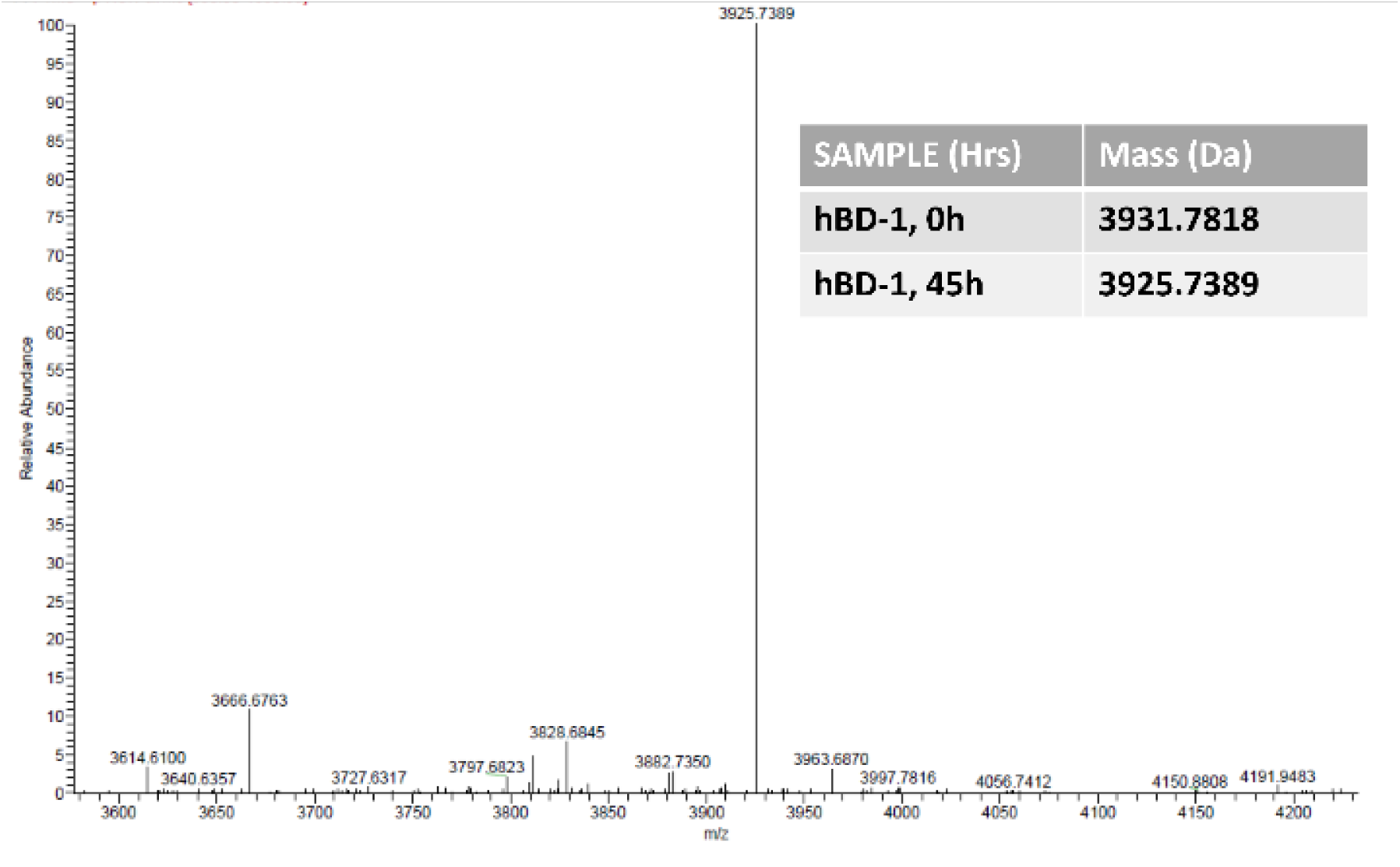
**Synthesis, Folding and Characterization of hBD-1 Using Controlled Oxidation Protocol and LC-MS Method, Yield >97 %, Purity >96%.**

### B: Cytotoxic Ppotential of hBD-1 in Cancer Cells

First, the antitumor potential of hBD-1 was tested in several prostate and triple negative breast cancer cells including on Fas resistant PC3 and TNBC MDA-MB-231 cells. Cancer cells were plated in 12 well plates 24 hrs before to a 60 % confluence were treated with various concentrations of hBD-1 for further 24-72 hrs and were analyzed by FACS and fluorescent microscopy with Calcein A/EtBr–dimer live/dead cells staining kit (Molecular probes). Fig. 2A shows concentration dependent cell death induced by hBD-1 in representative PC3 cells with IC_50_ around 20-30 μM (N =3, P <0.005). However, neither normal prostate epithelial cells ABC-TC3995 nor cardiomyocytes (target for doxorubicin or docetaxel toxicity) did not show significant change in apoptosis profile when treated with hBD-1 even up to 250 μM. (Fig. 2 B). hBD-1 was also combined with the standard first line chemotherapy treatments such as Doxorubicin or Docetaxel to see if they synergize the cell death. The results show that IC-50 combination of hBD-1 with either Doxorubicin or Docetaxel yielded IC-50 values by an order of magnitude compared to individual drugs (Fig 2B).

**Fig 2:**
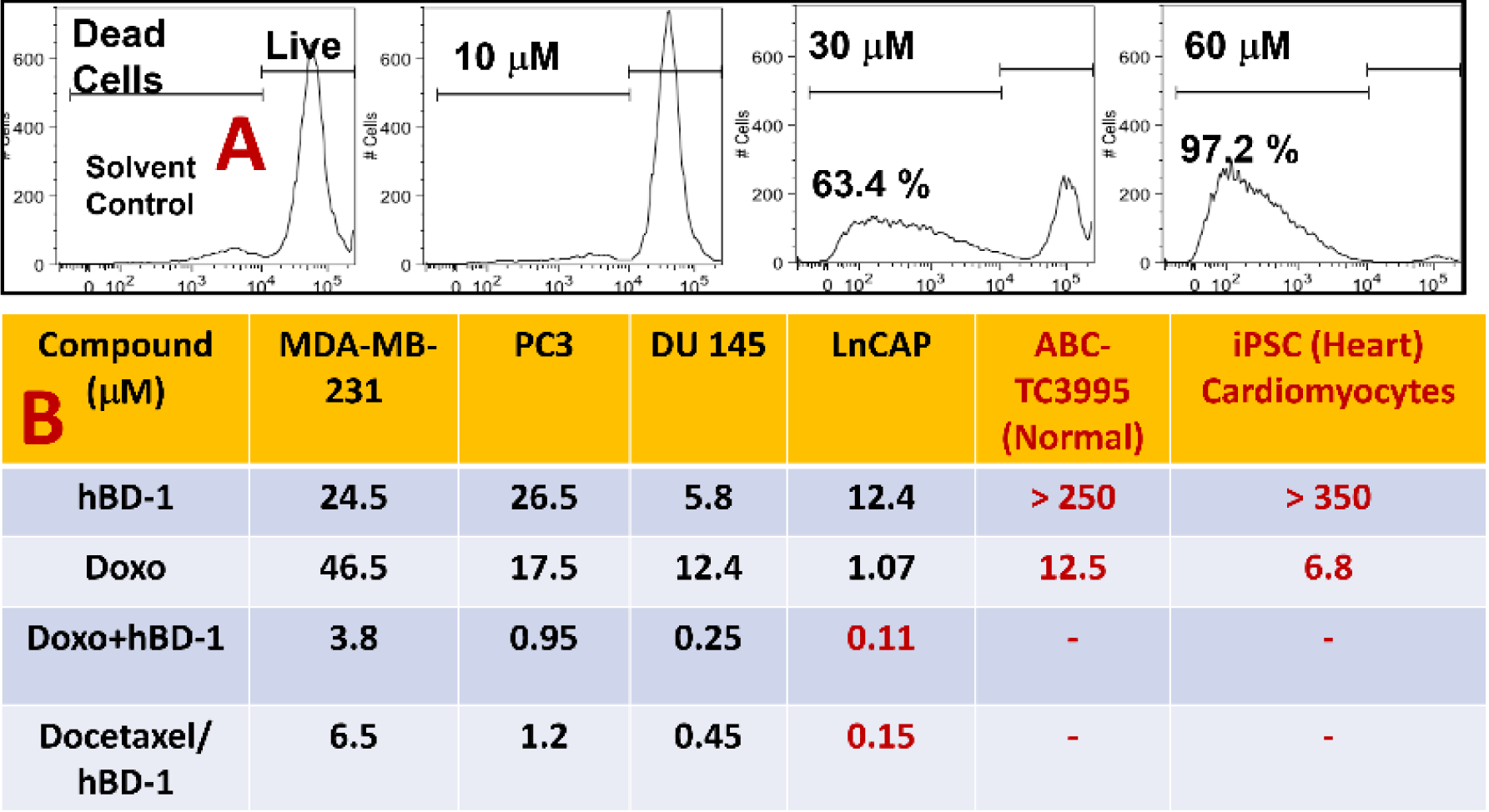
A Representative Prostate cancer cell PC3-FACS, B: Table 1: Relative Average IC-50 Values ( N=4, p < 0.005) for hBD-1, Doxorubicin and combination of hBD-l/Doxo/Docetaxel. TNBC Cells: MDA-MB-231, PC3, DU-145, LnCAP: Prostate cancer cells, Normal prostate epithelial Cells: ABC-TC3995 & Induced Pluripotent Stem Cells (iPSC Cardiomyocytes).

### C: Synergistic Potential of hBD-1 with Doxorubicin

The hhypothesis of sensitizing tumor cells in order to evoke a better response from chemotherapy was verified by incubating Fas resistant, MDA-MB-231 cells with 25 μM hBD-1 for 12 hrs before administering different concentrations of Doxorubicin (1, 2 and 3 μg/mL). Increase in cell death was assessed by fluorescent microscopy after 24-72 hrs. There is a dramatic effect of increase in cell death induced by hBD-1 in cells compared to the cells treated with only Doxorubicin (Fig. 3 compare C-D-E with F-G-H respectively). The pretreatment of MDA-Mb-231 cells with hBD-1 showed more than 90% cell death compared to either hBD-1 conjugate or Doxorubicin alone. These data indicate that hBD-1 sensitizes MDA-MB-231 cells to evoke a better response from front-line treatment Doxorubicin. Similar results are shown for prostate cancer PC3 cells where synergy with Doxorubicin was established (Supplementary Materials, Fig 1). The combination effect is revealed in the drastic reduction of IC-50 for the combination of hBD-1 with Doxorubicin and docetaxel (Table 1, 26.5 μM for hBD-1 Vs 0.95 μM and 1.2 μM respectively.

**Fig 3:**
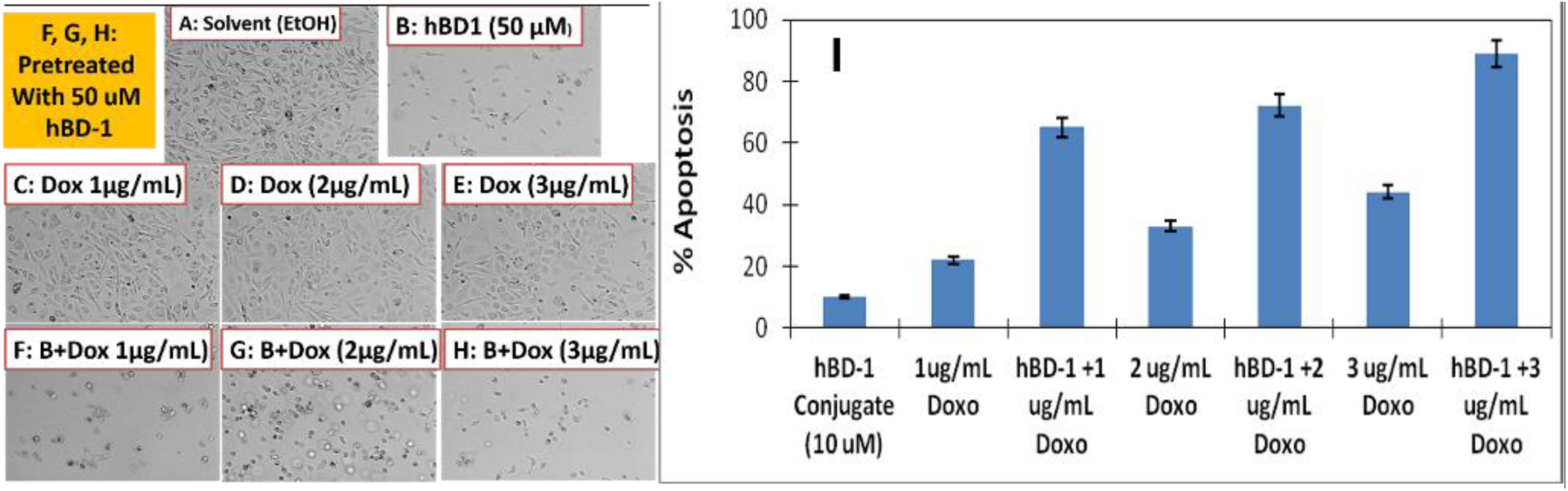
Treatment of MDA-MB-31 TNBC cells with hBD-1 resulted in cell death while in combination with Doxorubicin there is a synergistic cell death: Photomicrographs of cancer cells, A: Solvent only, B: 50 mM hBD-1 alone, C,D & E: Doxorubicin alone. F-H in combination with 50 mM hBD-1, I: Bar graph, N =5, p < 0.003

### D: Activation of Cell Death Pathway CD95 to Sensitize Tumor Cells

CD95 cell death pathway plays an important role in triggering apoptosis. Tumor sensitivity to chemotherapy *in vivo* is shown to be dependent on spontaneous baseline tumor apoptosis index^5f^ created by the deficient activation of apoptosis pathways. Fas resistant TNBC (MDA-MB-231, MMP2 positive, thioredoxin positive) cells were pretreated with hBD-1 conjugate for 6 hrs. after which cells were treated with anti-Fas antibody (ZB4, Enzo life sciences, Farmingdale, NY) or control antibody Mouse-isoform-IgG (Rockland Immunochemical, Gilbertsville, PA) and assessed for CD95 expression by FACS analysis. At 24 hrs. after incubation with 10 μM of hBD-1 conjugate, the cells stained for CD95 expression by FACS showed a log order change in the CD95 expression compared to control cells (Fig 4). This result indicates a potential mechanism of apoptosis activation by hBD-1 conjugate.

**Fig 4:**
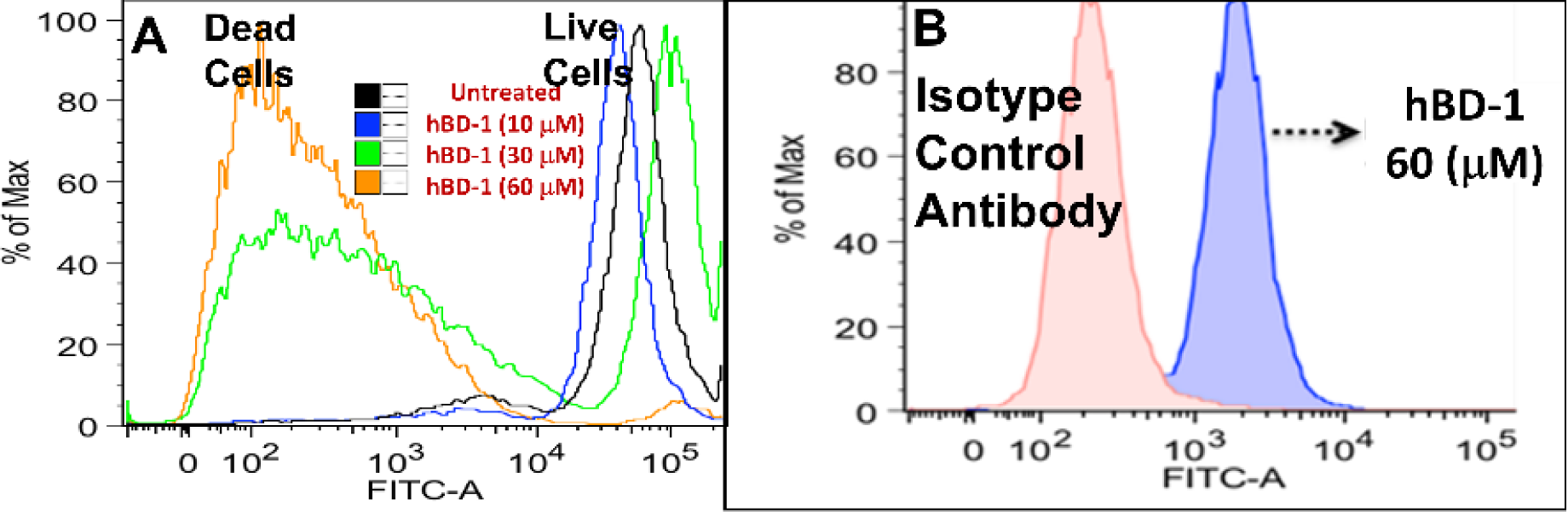
A: Dose dependent CD95 expression after hBD-1 treatment (10, 30 and 60 μM) in MDA-MB-231 cancer cells stained with FITC-anti-CD95 antibody n =3, p < 0.004, **B:** FACS analysis in MDA-MB-231 cancer cells treated with Isotope matched control antibody Vs hBD-1, X axis: Cells with fluorescence intensity, Y axis: Number of cells with specific fluorescence intensity, **Note:** Live cells tend to have intact unfragmented DNA show strong staining compared to apoptotic cells with fragmented DNA which show diffused and low intensity staining pattern.

E: Oxidation of Thioredoxin (Trx) by hBD-1: Recent investigations revealed that hBD-1 under reduced conditions possess better biological activity than in the oxidized form. Since Trx is a validated tumor specific biomarker and involved in deactivating ASK1 pathway, we decided to investigate the interaction between hBD-1 and Trx. Fig 5 shows concentration dependent treatment of hBD-1 with Trx where the intensity of the reduced form of Trx band at 14 KDa reduces at the expense of oxidized version at 28 KDa band. The data also showed positive control for hydrogen peroxide (H_2_O_2_, 2 mM) and standard Trx inhibitor PX-12 which is known for Trx inhibition via oxidation mechanism. We corroborated the results by incubating Trx with hBD-1 and followed the reduction of hBD-1 to linear form using LC-MS. The oxidized form of hBD-1 showed R = 6.9 min (corresponding to MS peak at 3025 M+) at t = 0 while after 2 hrs, the new peak appeared at Rt = 5.9 min which corresponds to M+ ion 3031 which coincided with the reduced linear form of hBD-1.

**Fig 5:**
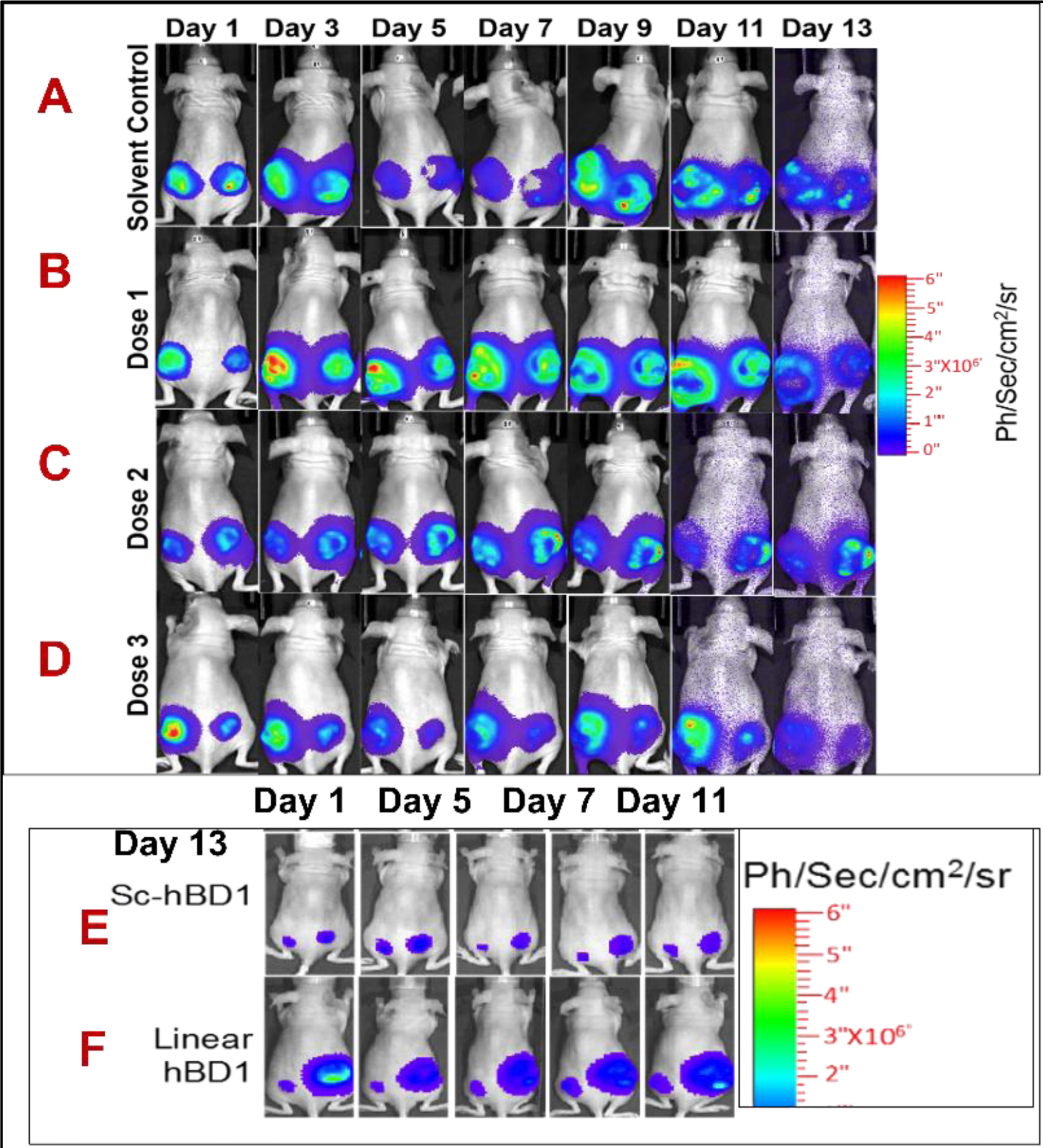
Dose dependent Tumor Regression by hBD-1 in TNBC MDA-MB-231 Tumor Xenograft by hBD-1 at Day 1-13. A) Untreated solvent control, B) 50 mg/Kg BW Dose 1, C) 100 mg/Kg BW, Dose 2, D) 200 mg/Kg BW, E) Scrambled hBD-1 (200mg.Kg, BW0 and F) Linear hBD-1 (200mg/Kg, BW, n = 5, p , 0.004.

### E: Anti-Tumorigenic Potential of hBD-1

Therapeutic evaluation of hBD-1 was carried out using MDA-MB-231 tumor xenograft in female nude mice with age group of 4 to 5 weeks. Nude mice were implanted with 10 million MDA-MB-231 cells stably expressing FLuc-EGFP fusion protein on either flank regions of hind limps to allow tumor growth. Tumor xenografts with a minimum size of 50 to 100 mm^3^ were selected for optical imaging studies. To perform therapeutic evaluation, nude mice with tumor xenograft were divided in to 5 groups comprising 5 animals in each group. One set of experiments consist of a) untreated control , b) 50 mg/Kg BW dose 1, c) 100mg/Kg BW, dose 2 and d) 200mg/Kg BW, dose 3. Another set of experiments consists of two separate controls a) scrambled hBD-1 and linear hBD-1 without folding. Bioluminescent signals were captured in animals from all 5 groups just before treating with hBD-1. Animals were treated with vehicle control (250µL physiological saline containing 10% PEG400), and other 2 control groups were treated with 200 mg/Kg BW hBD1 of scrambled hBD-1 and linear hBD-1 in 250µL physiological saline containing 10% PEG400. hBD-1 was administered by intra-peritoneal route In parallel to bioluminescence signal acquisition, tumor volume was measured using standard calipers in animals of all five groups every alternative day Fig 5 A shows that Tumor volume in untreated group was continuously growing while, the maximum dose (Fig 5D, 200mg/Kg BW) showed a significant tumor regression. It is noted that the tumor volumes at lower dose 50mg/Kg BW is similar to untreated control while medium dose 100mg/Kg has relatively less impact on tumor regression and tumor size was not growing as fast as the untreated control group. Unfortunately even the highest dose kept the tumor regression not below the initial tumor volume, but the tumor burden is significantly less compared to untreated control. The graphic representation of the tumor regression is shown in Fig 6.

**Fig 6:**
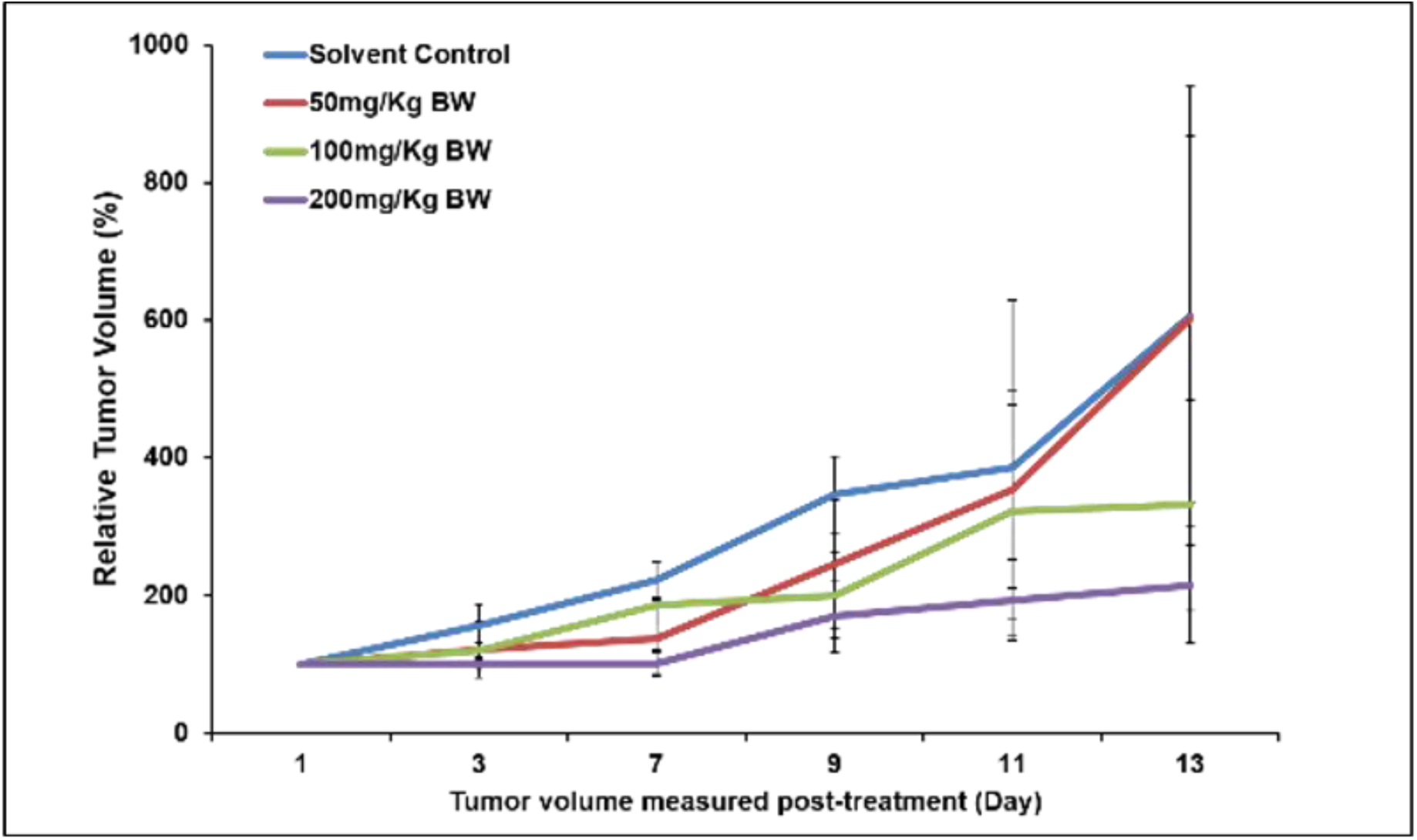
Graph showing average tumor volume measured every alternate day using vernier caliper. Tumor volume was analyzed and recorded as mm3 with final results presented as a relative tumor volume in percentage compared with initial tumor volume n = 5, P < 0.05.

### F: Histology of Tumor (target) vs Non-Target Organ Tissues

At the end of experimental period, animals from all 5 groups were euthanized and tissues samples from tumor, liver, kidney, heart, and spleen were fixed in 4% paraformaldehyde in PBS for histological and toxicological examinations.

## Discussion

Human Beta Defensin-1 is a tetra sulfide array with S-S folding which is known for antimicrobial, antifungal and to some extent anti-tumor potency. Defensins, in general are key effector molecules of innate immunity that protect the host from infectious microbes. Different versions of hBDs (hBD-1, hBD-2 and hBD-3) carry different kinds of biological activities, while hBD-1 is shown to be a potential tumor suppressor. Human Beta Defenisn (hBD-1) is one such immunomodulatory peptide which is lost at high frequencies in <u>malignant</u> prostate, renal and triple negative breast cancers (82 %, 65 % and 90 % respectively), while high levels of expression are maintained in benign regions^5,16^. hBD-1 is a tumor suppressor^16^ and its cancer specific loss was due to the loss of tumor suppressor gene located in chromosome 8p23 and to metastasis^16^. Delivering hBD-1 *in vitro* and *in vivo*, or its reexpression in malignant cells or overexpression of hBD-1 gene in cancer cells resulted in apoptosis^17^. Reexpression of hBD-1 through knock down of PAX2 oncogene promoted cancer cell death by 45 % compared to control. In essence, immunity protein hBD-1 role is to protect cells from becoming cancerous in the first place, its loss may be linked to onset of cancer and hence, overall restoration of hBD-1 back to the basal level promoted cancer cell death. The loss of expression of this protein by transformed epithelium may therefore inhibit host immune recognition or defensin-mediated tumor cell cytotoxicity. Given the proof of concept on the hypothesis of restoration of the lost hBD-1 protein, it may be possible to predict the early onstage of cancers, particularly for veterans who likely to lose their age-related immunity. This indicates that hBD-1 loss could be a potential biomarker for the onset of cancer which may lead to the use of hBD-1 expression as a valuable diagnostic tool to prevent the onset or to monitor cancer progression.

Current treatments for cancer, in general, are cocktails of many chemotherapeutics targeting different biological targets. However, it is uncommon that the drugs are targeted to dysregulated pathways. Amongst many survival pathways for cancer cells, downregulation of CD95 cell death pathway and inhibition of ASK1 pathway are relevant to hBD-1 treatment. FACS analysis of Fas resistant TNBC MDA-MB-231 cells.

Equally important in the failure of current treatments is the cancer cells ability to bypass intervention. Added to the immunity compromise through the loss of hBD-1, the cancer cells ability to deactivate cell death pathways (e.g. CD95, ASK1) resulted in desensitizing cancer cells to intervention, irrespective of the nature of intervention^2^. Cancer cells including TNBC overexpress thioredoxin (Trx) which inhibits activation of cell death pathway ASK1 and reduces the response from, for example cis-platin, mitomycin C, doxorubicin^21^ making them ineffective. Such tactics trigger high dose treatment, which in turn increases side effects, development of resistance, stopping the treatments temporarily which allows cancer cells to rejuvenate themselves. New rationale involving immunity modulation combined with sensitizing cancer cells may trigger significant clinical benefits compared to the existing standard of care (SOC) treatments.

Triple negative breast cancer (TNBC) is a devastating disease which disproportionately affects African Americans compared to Caucasians, accounting for 50 % death with a poor prognosis. The mortality rate for TNBC patients is 40% within 5 years of diagnosis^34^. Patient survival after cancer recurrence rarely extends beyond 12 months^35^. Unfortunately, no targeted receptor based FDA approved treatments (e.g. Tamoxifen, Herceptin etc.) work since TNBC patients do not carry biomarkers (e.g.; ER, PR and HER2 negative) for which the drugs are targeted. That compels chemotherapy as a first line of treatment for TNBC, despite a) high off-target toxicity, b) induction of stemness to expand cancer stem cells, c) elimination of bone marrow cells and may have to be stopped due to lymphedema^39^. This is a serious clinical problem which results in an increase in recurrence rate (13% for kidney cancer, 36% for breast and almost 100% for brain cancer) and makes tumors refractory to future treatments^40^. Defective apoptosis machinery combined with loss of immunity play major role in the resistance development^42^. For chemotherapy to be effective, apoptosis machinery has to be intact or not dysregulated. Without effective treatment strategies, roughly 20,000 of TNBC patients die yearly in the U.S. alone, making the development of potent TNBC therapies an unmet and an urgent medical need^36^.

The basic premise of our project is to combine the restoration of lost hBD-1 and activating cell death pathways simultaneously as a strategy to sensitize tumor cells with low or non-responsive to chemotherapy. Improving the clinical performance of the existing chemotherapeutics by making them work at lower doses, thus expanding the therapeutic indices. Such an approach may permit to reduce the dose related off-targeting adverse side effects of chemotherapy. Given the safety and efficacy of hBD-1, physicians may have large window of dose manipulations of combination therapy to tune dose regimen to optimize the treatment plans for patients. Accordingly, the anti-tumoregenic potential of hBD-1 was tested in a number of cancer cells which showed reasonable IC-50 (in early micromolar range) values which are similar to manty chemotherapeutics. However, hBD-1 being an endogenous protein seems to have minimum cell death potential for normal cells including breast epithelial cells (MCF-10A), cardiomyocytes iPSC (induced pluripotent stem cardiomyocytes) and bone marrow cells (MCF-001A). These are the cells, which in general are affected by chemotherapy at high doses. Reducing the chemotherapeutic dose without compromising on efficacy is expected to reduce cell death in these normal cells.

Downregulation of CD95 pathway by cancer cells is well correlated to the insensitivity of tumor cells to chemotherapy which means higher doses may be required to induce apoptosis in resistant tumor cells which in turn leads to high side effects. Dose dependent hBD-1 treatment in MDA-MB-231 cells using FACS reveals a log order of increase in fluorescence intensity of the apoptotic dead cells indicated by FITC Anti-CD95 antibody staining compared to an isotype matched control antibody (Fig 4A-B respectively). On the other hand, scrambled hBD-1 or linear hBD-1 controls did not show the change in the intensity of fluorescence peaks post treatment.

Thioredoxin (Trx) is a tumor specific protein overexpressed in many cancers including TNBC. Thioredoxin (Trx) is a small redox-regulating protein, which plays a crucial role in maintaining cellular redox homeostasis and cell survival^22^. The tumor environment is usually under either oxidative or hypoxic stress and both stresses are known to up-regulate Trx expression. Apoptosis signal-regulating kinase 1 (ASK1), a mitogen-activated protein kinase, plays a key role in the pathogenesis and resistance development to chemotherapeutics in cancer^23^. Trx inhibits ASK1 kinase activity by direct binding to its N-terminal coiled-coil domain (NCC). The free SH groups carried by Trx could play a potential role in the biological activity of Trx, particularly on the binding with ASK1 protein. We along with others have shown that poly sulfides can interact strongly with Trx. We have reported recently that plant defensin MsDef1 with tetra sulfide bond array similar to hBD-1 oxidized Trx in TNBC cells and activated ASK1 pathway. hBD-1 is shown to be benign under normoxic conditions while, under hypoxic conditions (e.g. tumor), reduction of cysteine-cysteine (S-S) bonds by thioredoxin unmasks biological activity^24^. hBD-1 oxidizes thioredoxin (see HPLC results Fig 2 in ref 19 and our results) and deactivates Trx biological activity. Thioredoxin is a well-studied clinical target which is known to lower the response rate to docetaxel, cis-platin and doxorubicin^21^. Alteration of intracellular tumor Trx from reduced form to oxidized form *in vivo* by Trx inhibitors sensitized tumor cells to drug induced apoptosis^21.^ Our data on oxidation of Trx by hBD-1 by both Western Blotting and RP-LC-MS is likely to release ASK1 protein binding from Trx, thus activating ASK1 cell death pathway which may enhance the sensitivity of tumor cells for chemotherapy. The relevance of the more potent reduced form of hBD-1 by Trx was demonstrated by Schroeder et.al who showed that immunohistochemical staining of hBD-1 reduction is catalyzed by Trx in human epithelia. More work is needed to show the oxidation of Trx by hBD-1 in vitro using cancer cells and the quantification of the phosphorylation of the specific Thr485 residues of Trx as we have shown in plant defensin MsDef1 which is very similar to hBD-1.

Dysregulation of cell death pathways is a major culprit in making cancer cells desensitize to interventions. By activating these specific cell death pathways it is expected that tumor cells get sensitized to chemotherapy. Hypothesis of sensitizing tumor cells in order to evoke a better response from chemotherapy was verified by incubating Fas resistant, MDA-MB-231 cells with different doses of hBD-1 for 12 hrs before administering different concentrations of doxorubicin (1, 2 and 3 μg/mL). Increase in cell death was assessed by fluorescent microscopy after 24-72 hrs. There is a dramatic effect of increase in cell death induced by hBD-1 in cells compared to the cells treated with only doxorubicin (in Fig. 3 compare C-D-E with F-G-H respectively). The pretreatment of MDA-MB-231 cells with hBD-1 showed more than 80% cell death compared to either hBD-1 conjugate or doxorubicin alone. These data clearly indicate that hBD-1 sensitizes cancer cells to evoke a better response from front-line drug doxorubicin. Similar results are shown for docetaxel in prostate cancer DU-145 cells which is also one of the front-line treatments for prostate cancer (See supplementary materials). The combination effect is revealed in the drastic reduction of IC-50 for the combination of hBD-1 with doxorubicin and docetaxel (see Table 1, 3.8 μM for hBD-1 Vs 45 μM for Doxorubicin respectively). The synergy of hBD-1 and Doxorubicin was also assessed through combination index calculated using Towley method(CI) which showed less than 1.00 for several dose combinations (see Supplementary materials).

Although exogenous administration of hBD-1 has reasonable anti-tumorigenic properties it is uncertain how it behaves in vivo. For *in vivo* therapeutic evaluation, we used tumor xenograft of MDA-MB-231 stably expressing FLuc-EGFP fusion protein. Nude mice were implanted with 10 million MDA-MB-231 cells stably expressing FLuc-EGFP fusion protein, on either flank regions of hind limps. Fig 6 showed a dose dependent tumor regression from 50 mg/Kg to 200 mg/Kg compared to untreated control (Fig 7, tumor volume is reduced from 580 mm^3^ to 150 mm^3^ for 200mg/Kg dose). There were no significant changes either in the bioluminescence signal or in the tumor volume for scrambled hBD-1 or linear hBD-1 signifying the specificity of hBD-1 attributed to full folding of the protein followed by presumably reduction in vivo. The differential IC50 values in cancer cells vs normal cells (Fig 1) corroborated well with the in vivo data. This is a significant result despite the dose is relatively high compared to standard chemotherapy. However, histological characterization of target tumor Vs non-target organs may throw more light on the safety of hBD-1 in vivo at the efficacy dose. Animals from all the groups were euthanized and tissue samples from kidney, liver, heart, spleen, and tumor were fixed with formalin and embedded in paraffin for histological and toxicological examinations. Tumor tissue sections from animals treated with hBD1 showed marked tissue damage in cellularity (**Fig 8 A**). On the contrary, histological analysis of non-target tissues (heart, kidney and liver) did not show significant damages compared to untreated control (**Fig 8 B, C & D).** We have reported, on the contrary, that Doxorubicin induced cardiotoxicity at the efficacy dose (21 mg/Kg) in vivo rat model. Since hBD-1 is a natural protein endogenously produced it is not expected to be toxic unless a very high dose is used.

The target to non-target ratio of the drug determines efficacy and toxicity. Based on the limited data, one can hypothesize that HBD-1 might target tumor specific intracellular Trx overexpressed by many cancers. The activation of both dysregulated CD95 and ASK1 pathways may be responsible for sensitizing low or no responsive tumor cells to chemotherapy. The selectivity of hBD-1 for cancer cells is based on the cell penetrating ability of cationic trisulfide folded hBD-1, the potential biological activity was unmasked by tumor specific Trx. The high cationicity of hBD-1 is due to the low pH environment of cancer cells which increases the cationicity from +4 to 7 due to protonation of several amino acid residues of hBD-1. Table 1 summarizes the selectivity of hBD-1to cancer.

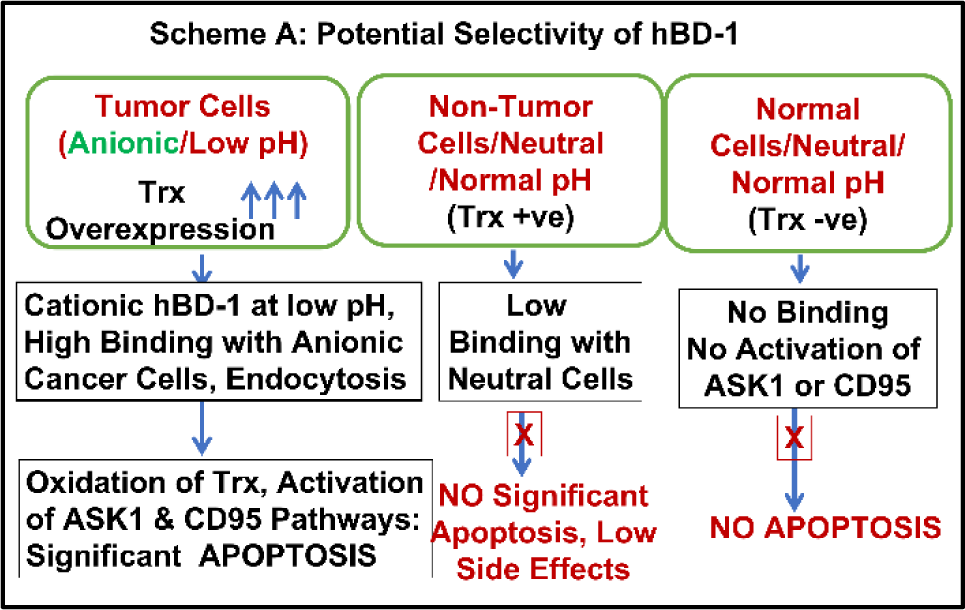

## Conclusions

Loss of natural immunomodulatory protein hBD-1 in many cancers combined with the inherent characteristics of cancer cells ability to dysregulate cell death pathways led to the hypothesis that whether restoration of the lost hBD-1 in combination with activation of cell death pathways a viable strategy for the treatment of cancer. Our approach is to use hBD-1 as a targeted tumor sensitizer to make nonresponsive tumor cells respond better to the current treatments, particularly for TNBC patients who depend on nonspecific chemotherapy. in The limitations of the study includes the detection of reduced form of hBD-1 as a biological entity in vivo , longer treatments time to see the tumor volume to be near zero and extension of synergy of Doxorubicin or any other front line chemotherapeutics to in vivo. However, the established antimicrobial properties of hBD-1 extension to anti-tumorigenic natural substrate may open a new way of using hBD-1 as a neoadjuvant to chemotherapy. The new mode of understanding the molecular biology of cancers in terms of dysregulated pathways and revamping them to make chemotherapy effective at lower dose may improve the safety profile of the current treatments significantly.

## Experimental

### A) Synthesis and Characterization of hBD-1

The linear hBD-1 was synthesized by Ambiopharma under GLP conditions. Both HPLC analysis and mass spectrum showed > 96 % purity and M+ ion at 3931.7 corresponding to the linear hBD-1 without cyclization of S-S bonds. 30 mg of the sample linear hBD-1 was dissolved to the concentration of 0.6 μg/μL in 50 mM phosphate buffer containing 1M Guanidine HCl at pH 7.5. The solution was sealed into a tube through which low pressure air bubbled continuously. The sample was aliquoted at different time points from 0 h to 45h. The samples were desalted with C18 zip tip and run on LTQ-Orbitrap Velos by direct infusion. The samples were run with high resolution (60,000, LTQ-Velos Pro Orbitrap LC-MS/MS). The final sample at 45h was desalted using Sepak SPE and the eluted sample was dried down. HPLC showed a purity of > 96 % while LC-MS showed a molecular ion peak at 3925.7 which corresponds to loss of exact six protons giving raise to folded final peptide. The correct folding of the protein was verified through circular dichroism (CD) and retention of biological activity in cancer cells.

### B) Viability Assessment and IC-50 Determination

Recently, we have described MTT assay for plant defensin, and we followed the same procedure. MDA-MB-231 cells were maintained in Dulbecco’s modified eagle medium (Sigma, USA). Breast normal epithelial cells MCF10A (American Type Culture Collection, Manassas, VA) were cultured in DMEM/Ham’s F-12 (GIBCO-Invitrogen, Carlsbad, CA) supplemented with 100 ng/ml cholera toxin, 20 ng/ml epidermal growth factor (EGF), 0.01 mg/ml insulin, 500 ng/ml hydrocortisone, and 5% chelex-treated horse serum. The cells were cultured at 37 °C in a 90% humidified incubator with 5% CO_2_. When the cells were 80% confluent, they were sub-cultured to a fresh media. Cardiomyocytes were derived from this engineered stem cell clone line as follows. Stem cell aggregates were formed from single cells and cultured in suspension in medium containing zebrafish bFGF (basic fibroblast growth factor) and fetal bovine serum. Upon observation of beating cardiac aggregates, cultures were subjected to blasticidin selection at 25 ug/ml to enrich the cardiomyocyte population. Cardiomyocyte aggregate cultures were maintained in Dulbecco’s modified Eagle’s medium (DMEM) containing 10% fetal bovine serum during cardiomyocyte selection through the duration of the culture prior to cryopreservation. At 30 to 32 days of culture the enriched, stem cell-derived cardiomyocytes were subjected to enzymatic dissociation using 0.5% trypsin to obtain single cell suspensions of purified cardiomyocytes, which were *>*98% cardiac troponin-T (cTNT) positive. These cells (iCell1Cardiomyocytes) were cryopreserved and stored in liquid nitrogen before delivery to Ionic Transport Assays from Cellular Dynamics International, Madison, WI.

## Combination Index Calculations

Combination Indices were calculated using CompuSyn software, V. 1.0 (Biosoft, Cambridge, UK). Drug interactions were classified by determining a combination index (CI) recognized as the standard measure of combination effect based on the Chou-Talalay method. The CI values were obtained over a range of fractional cell kill levels (Fa) from 0.05 to 0.95 (5-95% cell kill). Based on the Chou-Talalay method(1,2,3,4), CI < 1 means synergism, CI = 1 means additivity, and CI > 1 is interpreted as antagonism.

For FACS analysis, we plated MDA-MB-231 cells in 6-well plates at 2 × 105 cells/well in DMEM medium supplemented with 10% FBS. We grew cells overnight and changed to DMEM medium supplemented with 10% FBS and exposed them to the required conditions (e.g. various drugs at different concentrations), each condition being represented by three wells; unexposed cells served as controls in every experiment. Cells were then allowed to grow for 6, 12, 24 and 48 h post exposure to drugs before collecting them for FACS analysis. We collected all cells by trypsinization and suspended them in 0.5 ml of ice -cold PBS. Briefly, cells were spun, washed once in PBS, and suspended in 0.5 ml PBS containing 1 μl BFC. After 30 min of incubation at room temperature, we washed the cells once with PBS, re-suspended in 0.3 ml PBS, and used for FACS analysis. Cells were FACS analyzed using a Guava-FACS analyzer [EMD Millipore], and we analyzed the generated data using FlowJo 8.6.6 Software [Tree Star].

### D). Western Blotting

The Trx Western blotting was performed as described previously, with minor modifications^24^. Briefly, 3 x 10^6^ cells were lysed in G-lysis buffer (50 mM Tris HCl, pH 8.3, 3 mM EDTA, 6 M guanidine–HCl, 0.5% Triton X-100) containing 50 mM iodoacetic acid (IAA; pH 8.3). For each experiment, control plates, for identifying Trx redox state bands in the Western blot, were also incubated with 2mM H_2_O_2_, for 10 min at room temperature, before incubation with 50 mM IAA. Subsequently, the lysates from all cells were incubated in the dark for 30 min with the IAA. The lysates were then centrifuged in G-25 micro spin columns (GE Healthcare). Protein was quantified from the eluent using the Bradford protein assay, as previously described^25^.

### E) Synergy of hBD-1 with Doxorubicin

MDA-MB-231 or PC3 tumor cells (ATCC) were cultured in 24-well plate for 18 hours. hBD-1 was dissolved in culture medium and sonicated for 2 min each and repeated for 3 times. After treated the cells with different concentrations of Doxorubicin HCl (1, 2 and 3 μg /mL, Bedford Lab, Bedford, OH), keeping hBD-1 constant dose (50 μg/mL) constant and combination of Doxorubicin and hBD-1, the cells were washed three times with PBS (pH 7.4) and incubated with Cy5.5 Annexin-V (BD Bioscience Pharmingen) according to the instructions of the manufacture. The cells were imaged using Nikon Eclipse TE-300 fluorescence microscope and counted under Ex/Em 620 - 680nm nm/ 700 - 750nm.

### F) In Vivo Xenograft Assay

For in vivo therapeutic evaluation, we used tumor xenograft of MDA-MB-231 stably expressing FLuc-EGFP fusion protein. Cells containing luciferase and EGFP fusion constructs were generated at Stanford University and screened by FACS as reported earlier [41]. Nude mice were implanted with 10 million MDA-MB-231 cells stably expressing FLuc-EGFP fusion protein, on either flank regions of hind limps. After the tumor is grown to a size range of 50 to 100 mm3, nude mice with tumor xenograft were divided in to 3 groups comprising 4 animals in each group, and 1 control group with 3 animals. Bioluminescent signals were captured in animals from all 4 groups just before treating with AMP-001. Group with 3 animals were treated with vehicle control (250μL physiological saline containing 10% PEG400), and other 3 groups were treated with 50 mg/Kg BW, 100 mg/Kg BW, and 200 mg/Kg BW AMP-001 in 250μL physiological saline containing 10% PEG400. AMP-001 was administered by intra-peritoneal route 7 times with an interval of 48h. For optical imaging, animals were intraperitoneally injected with 3 mg of D-Luciferin in 100 μl PBS, 5 to 10 minutes before signal acquisition. All mice were imaged with a cooled CCD camera (Spectral Lago; Spectral Instruments Imaging, Tucson, AZ), and photons emitted were collected and integrated for a period of 15 seconds for 20 acquisitions for FLuc. For optical imaging, animals were intraperitoneally injected with 3 mg of D-Luciferin in 100 μl PBS, 5 to 10 minutes before signal acquisition. All mice were imaged with a cooled CCD camera (Perkin Elmer, Akron, OH), and photons emitted were collected and integrated for a period of 15 seconds for 20 acquisitions for FLuc. Images were analyzed by Spectral Instruments Imaging Software (Spectral Instruments Imaging, Tucson, AZ). To quantify the number of emitted photons, regions of interest (ROI) were drawn over the area of the implanted cells, and the maximum photons per second per square centimeter per steradian (p/sec/cm2/sr) were recorded. All animal experiments were conducted under the guidance of the Administrative Panel on Laboratory Animal Care (APLAC), Stanford University. The Institute animal research committees at Stanford approved all animals handling and National Institutes of Health guidelines. APLAC specifically approved the current study on animals funded by NIH grant with NIH animal welfare assurance number A3213-0. All animals used for research at Stanford University are under the care of the comparative medicine program. The proposed procedures are in accordance with the Animal Welfare Act (PL99-158) and the *Guide to the Care and Use of Laboratory Animals*. All animals (Balb/c, nude, females) were purchased from Charles River laboratories, (Wilmington, MA).

## Immunohistochemical staining

The xenograft tumor slides were incubated with the following primary antibodies: anti-CD31 was purchased from ABclonal and anti-Ki67 from Cell Signaling Technology (USA). Anti-rabbit or anti-mouse peroxidase-conjugated secondary antibody (ABclonal) and diaminobenzidine colorimetric reagent solution purchased from Dako (Carpinteria, CA) were used. The staining processes were according to standard methods.

## Statistical analysis

All experiments were performed at least three times. Data are presented as means +/− SD. All statistical analyses were performed using GraphPad Prism 6.0 (GraphPad, SanDiego, CA). One-way ANONVA and Student’s t-test were applied to determine statistical significance. A value of p*<*0.05 was considered statistically significant.

## Acknowledgements

Dr. Raghu Pandurangi acknowledges the collaborative efforts from Professor Ramasamy Paulmurugan from Stanford University.

## Contributions by Authors

1. Dr. Raghu Pandurangi, Conceptualization, Interpretation of data, Writing original draft, design of experiments.
2. Dr. Ramasamy Paulmurugan, All biological in vitro and in vivo experiments design and manuscript editing.
3. Thillai V. Sekar, data collection.

