## Supplementary Figures for "Restoration of the Lost Human Beta Defensin (hBD-1) in Cancer as a Strategy to Improve the Efficacy of Chemotherapy"

### Supplementary Materials

Figure S1: Mass Spectrum of hBD-1

| Sample ID | Experimental exact mass (Da) |
|---|---|
| hBD1 0 h | 3931.7819 |
| hBD1 4 h | 3930.7658 |
| hBD1 18 h | 3928.7499 |
| hBD1 24 h | 3927.7441 |
| hBD1 42 h | 3925.7319 |

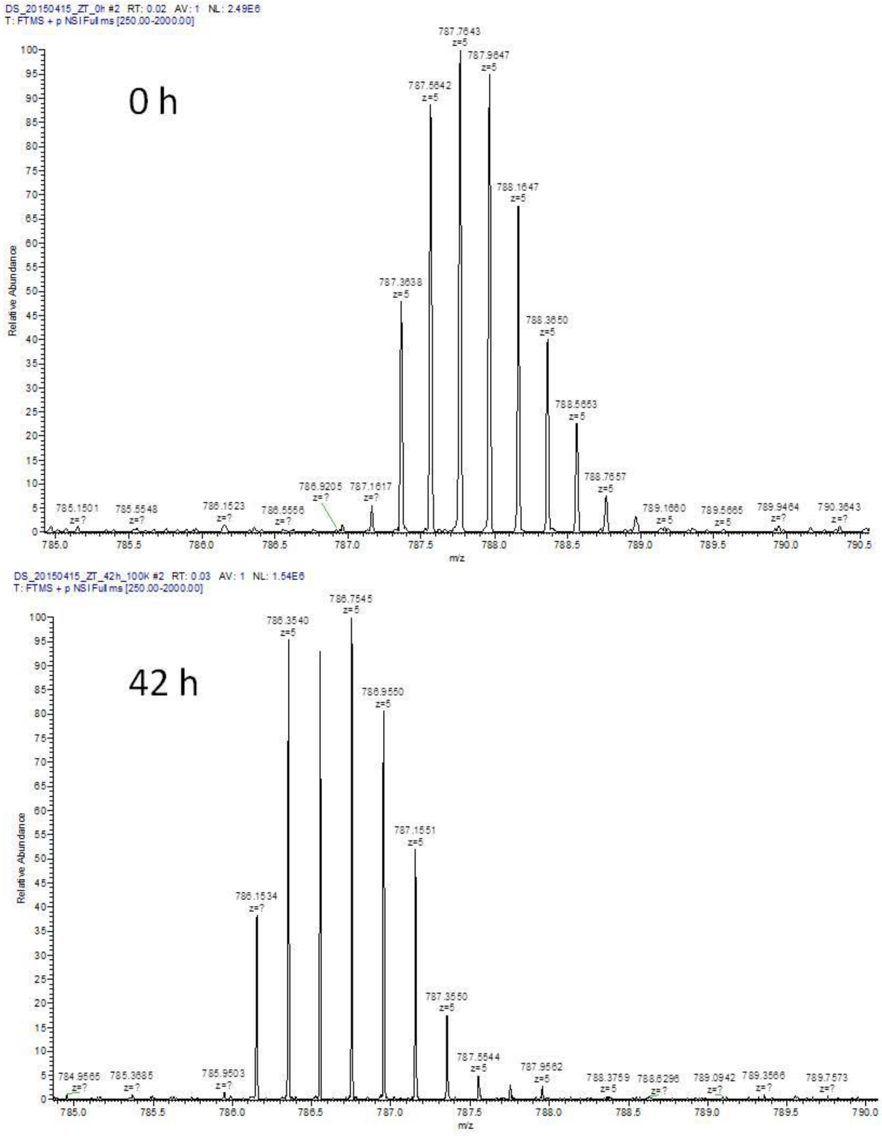

Figure S2: Mass Spectral Deconvolution at 0 hrs and at 42 hrs

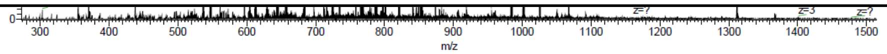


Mass Spectral Deconvolution of hBD1 at 0 h

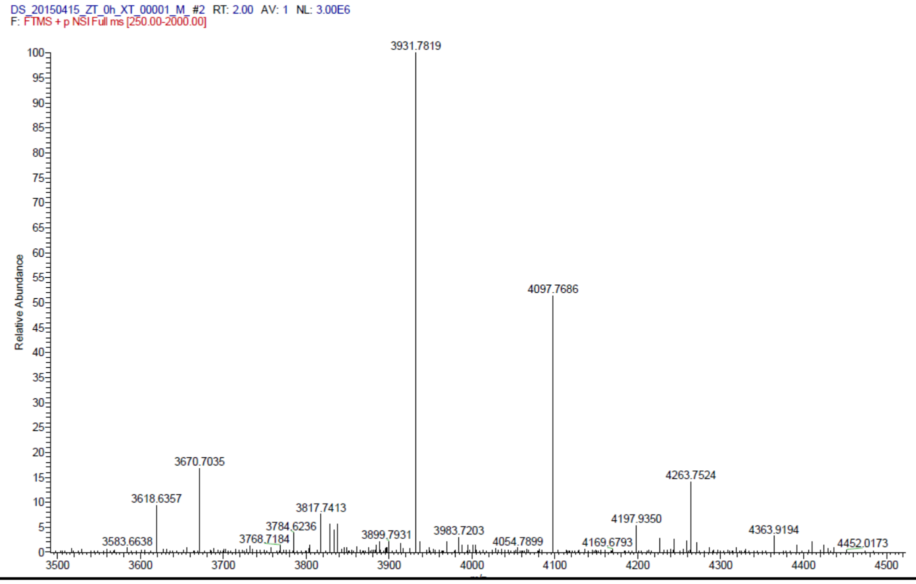


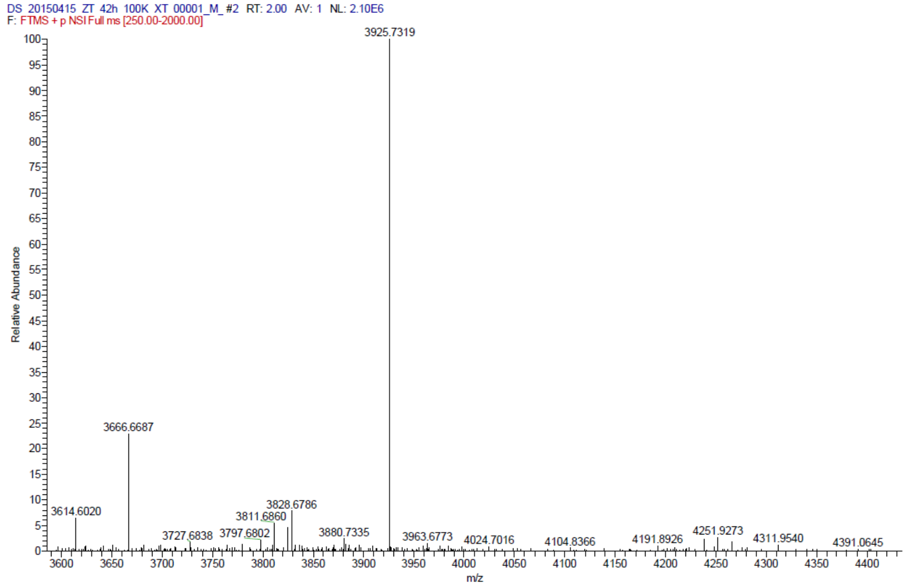
